# Assessing the Landscape of U.S. Postdoctoral Salaries

**DOI:** 10.1101/227694

**Authors:** Rodoniki Athanasiadou, Adriana Bankston, McKenzie Carlisle, Carrie Niziolek, Gary McDowell

## Abstract

**Purpose:** Postdocs make up a significant portion of the biomedical workforce. However, data about the postdoctoral position are generally scarce, including salary data. The purpose of this study was to request, obtain and interpret actual salaries, and the associated job titles, for postdocs at U.S. public institutions.

**Methodology:** Freedom of Information Act Requests were submitted to U.S. public institutions estimated to have at least 300 postdocs according to the National Science Foundation’s Survey of Graduate Students and Postdocs. Salaries and job titles of postdoctoral employees as of December 1st, 2016 were requested.

**Findings:** Salaries and job titles for over 13,000 postdocs at 52 public U.S. institutions and 1 private institution around the date of December 1st, 2016 were received, and individual postdoc names were also received for approximately 7,000 postdocs. This study shows evidence of gender-related salary discrepancies, a significant influence of job title description on postdoc salary, and a complex relationship between salaries and the level of institutional NIH funding.

**Value:** These results provide insights into the ability of institutions to collate actual payroll-type data related to their postdocs, highlighting difficulties faced in tracking, and reporting data on this population. Ultimately, these types of efforts, aimed at increasing transparency, may lead to improved tracking and support for postdocs at all U.S. institutions.

## Introduction

Postdocs make up a significant proportion of the academic workforce, and recommendations for reforms to improve the postdoctoral experience have been made for the last 50 years (Bankston and McDowell, 2018). In spite of this fact, the total number of postdocs in the U.S. is currently unknown (“ Biomedical Research Workforce Working Group: Biomedical Research Workforce Working Group Report. (Report to the Advisory Committee to the Director)”, 2012). Counting postdocs can be difficult due to the varying means by which they are administered, handled and classified among U.S. institutions, leading to the difficulty in keeping track of this population as a whole (Pickett et al., 2017; Schaller et al., 2017). As a result, very little data exist on specific aspects of the postdoctoral position itself, including salaries, benefits, fellowships, and career tracking.

Future of Research wants to champion, engage and empower early career scientists with evidence-based resources to improve the scientific research endeavor. It is therefore a stated goal to increase transparency around aspects of the postdoctoral position, and contribute to efforts in harmonizing the abundance of postdoc titles used within and across U.S. institutions (McDowell, 2016; Schaller et al., 2017). Given multiple titles, the term “postdoc” will be used in this publication to encompass all PhD-holding researchers who are not currently in faculty roles (either tenured, non-tenured or adjunct). This will inevitably include staff scientists, as this classification varies across institutions and is not consistently differentiated by job title from the nature of postdoctoral employment.

Very little open debate exists about the state of personal finances of academics. There is a widespread assumption in academia that postdocs are generally paid in accordance with the National Institutes of Health (NIH) NRSA scale (“Revised: Projected FY 2017 Stipend Levels for Postdoctoral Trainees and Fellows on Ruth L. Kirschstein National Research Service Awards (NRSA)”, 2016). However, these stipend levels only apply to postdocs on NIH training and fellowship funding mechanisms, such as T32 and F32 awards, and do not apply to all postdocs, including those on NIH research project grants such as R01s. As most postdocs are not funded through training and fellowship mechanisms, the salaries of a majority of postdocs in academia are the responsibility of institutions. Institutional policies exist to recommend salary minima and in practice many institutions use the NIH NRSA stipends as guidelines, according to the National Postdoctoral Association Institutional Policy Database and Report (National Postdoctoral Association, 2014). Although these guidelines are in place, the Fair Labor Standards Act (FLSA) essentially establishes the legal minimum for postdoc salaries (“United States Department of Labor: Final Rule - Overtime - Wage and Hour Division (WHD)”, 2016), and guarantees both minimum wage and overtime pay for full-time employees at 1.5 times the regular rate (“United States Department of Labor: Compliance Assistance - Wages and the Fair Labor Standards Act (FLSA) - Wage and Hour Division (WHD)”, 2016). The current legal minimum annual salary for full-time postdocs (with the exception of those tracking hours and being paid overtime) is therefore $23,660.

Updates to the FLSA were proposed in 2016 to raise the threshold annual salary for overtime exemption to $47,476. The proposal to update the FLSA emerged from a memorandum issued by President Barack Obama in March 2014 to “modernize and streamline the existing overtime regulations (Obama, 2014).” The proposed updates were announced by the U.S. Department of Labor (DOL) in May 2016 following a public consultation period, including raising the threshold salary for overtime exemption to $47,476, and automatically updating the threshold salary every 3 years. The FLSA updates were due to come into effect on December 1st, 2016. In response to these updates, previous work analyzed the compliance of 340 U.S. institutions with the FLSA ruling (Bankston and McDowell, 2016). However, an injunction granted nationwide on November 22nd, 2016 (“Anon: State of Nevada et al v. United States Department of Labor et al.”, 2016) halted implementation of the updates. In spite of this fact, a number of institutions maintained their commitment to raise postdoc salaries, and the NIH also moved forward with increasing NRSA stipends (“National Institutes of Health: NOT-OD-17-003: Ruth L. Kirschstein National Research Service Awards (NRSA) Postdoctoral Stipends, Training Related Expenses, Institutional Allowance, and Tuition/Fees Effective for Fiscal Year 2017”, n.d.). Some institutions, conversely, cancelled the previously-planned salary raises following the injunction (for a list of institutional policies after the injunction, see (Future of Research, 2016)). As the FLSA updates never came to pass, the mandate to raise salaries was therefore removed (Bankston and McDowell, 2016).

This change in the landscape of institutional postdoc policies, in addition to the difficulties of obtaining accurate counts of postdocs from institutions (Pickett et al., 2017), likely indicates that postdoc salary policies may not be effectively enforced across institutions. Therefore, the current study sought to evaluate individual postdoc salaries in the U.S. in a standardized fashion, by requesting and interpreting salary information using Freedom of Information Act (FOIA) requests. In order to provide an estimate of the number of postdoc salaries expected to be received from an institution in the current study, this was compared to the number of postdocs reported to be employed at that institution in health, science and engineering fields using data from the 2015 National Science Foundation’s (NSF) Survey of Graduate Students and Postdocs (GSS) (“National Science Foundation: Survey of Graduate Students and Postdoctorates in Science and Engineering”, 2015). While a true comparison is difficult because of reporting discrepancies in postdoc numbers (Pickett et al., 2017) and the exclusion of non-science, engineering and health postdocs from this data, this number served as a guide for the number of salaries expected from an institution.

This report describes the data collected and the implications of obtaining publicly available postdoc salaries as of December 1st, 2016. We analyzed postdoc salary data with respect to geographic region, gender, postdoc title and institutional NIH funding level. Although the breadth of our analysis is limited by the number of variables provided across universities, as well as institutional variability in being able to report actual annual salaries, the large sample size of the dataset allows us to draw meaningful conclusions. We find evidence of gender-related salary discrepancies, a significant influence of job title description on postdoc salary, and a complex relationship between salaries and the level of institutional NIH funding.

As a whole, this dataset allows us to better understand the U.S. postdoctoral workforce and inform data-driven strategies for future work.

We have gathered this information in an online resource (“Investigating Postdoc Salaries | Future of Research”, n.d.; Woolston, 2017). This work sheds light on the difficulty of reporting actual postdoc counts and salaries across U.S. institutions. It is possible that general postdoc conditions could be inferred through institutional policies (e.g. general salary information can be deduced through institutional policies on minimum salary; general benefits information can be concluded based on the number of postdocs classified into a certain benefits-offering job code). However, there are likely many factors that contribute to determining postdoc salary beyond institutional policies. Such factors may include: the postdoc’s previous experience; availability of laboratory funding; salaries of postdocs within the same lab or department; and subjective decisions about what a postdoc “should” be paid which are typically not verified against institutional policies. As postdocs tend to be hired by individual Principal Investigators, and not by the institutions as a whole, variability in salaries may exist within an institution in the absence of direct oversight. These differences in dealing with individual postdoc salaries within institutions contribute to the difficulty in discerning the factors that affect the actual salary received. We further suggest postdoc numbers and salaries could be used as a model to illustrate problems and recommendations for reporting on various aspects of the postdoctoral position. This is in line with our overall mission of increasing transparency about the scientific enterprise to empower early career researchers to make rational decisions about their career paths and using their passion for science to benefit society.

## Methods

### Data Collection

Postdoc salary data were collected using Freedom of Information Act (FOIA) requests made to 51 public universities with full-time postdocs. Requests were made for salaries and job titles of all full-time PhD- or MD-holding employees in postdoctoral research roles regardless of discipline. This data collection encompassed all public U.S. institutions with more than 300 science, engineering and health postdocs in the National Science Foundation’s 2015 Survey of Graduate Students and Postdocs (GSS) (“National Science Foundation: Survey of Graduate Students and Postdoctorates in Science and Engineering”, 2015). University systems with institutions containing fewer than 300 postdocs were also included in the requests (e.g. requests were made to each campus in the State University of New York system, regardless of size). While no FOIA-like mechanism exists for requesting data from private institutions, we obtained salary data from Boston University for postdocs supported on Research Project Grants/non-fellowship mechanisms. This data was volunteered, and is the only data from a private institution in this dataset. All FOIA requests asked for the following information:

*“The annual salaries, on December 1st 2016, of all full-time PhD or MD-holding postdoctoral employees and fellows, and employees under any other titles that encompass postdoctoral research roles, at [INSTITUTION NAME]”*.

Letters also asked for a written explanation of any denial of all or a portion of the FOIA request. Requests were only denied in two instances. The first is University of Utah, which denied the request on the grounds of the effort required, but was later required to provide the data on appeal to General Counsel. The second is The University of California Public Records Office, which rejected the request across all campuses similarly citing the effort required and that the fact that this data was not available. However, the University of California Santa Barbara had already provided the requested data when this claim was made.

Data from all 52 institutions were collated (“Investigating Postdoc Salaries | Future of Research”, n.d.) and represent salary information provided for 13,068 postdoctoral scientists in the U.S. around the requested date of December 1st, 2016. There is some margin of error around the date as some institutions explicitly stated that data was from slightly before or after this date. Also there are individual instances (for example, the University of Illinois) where we can say with certainty that the number of postdoc salaries received is incorrect, and where data received had reporting errors, most significantly in the instances where we did not receive annual salaries, but instead the amount of compensation that passed through payroll, leading to some of the extremely low salary amounts reported. Due to technical reasons, the UT San Antonio was only included in certain analyses as indicated in the main text (51 university dataset).

Responses were supplied in both electronic and paper format, with paper-based records being hand-entered in the database. These data are not a result of postdocs themselves reporting salaries in a survey (i.e. self-reporting). Because of the FOIA mechanism employed, universities likely retrieved data from a payroll or HR database in a more automatic fashion, which may or may not be the same method used when departments are reporting data for similar efforts on postdocs such as the NSF’s GSS (Pickett et al., 2017). In cases where salaries were reported for less than 100% effort (82 postdocs in 5 institutions), the annualized salary for 100% effort was calculated and used in the analyses.

### Data Preprocessing and Analysis

#### Gender assignments

For the postdocs whose names were available, gender was inferred from, and assigned to, the data entries based on each postdoc’s name using the Ethnea algorithm (Torvik and Agarwal, 2016) using the geographical distribution of the postdoc (i.e. author)’s affiliations in PubMed over time to predict ethnicity and apply an ethnicity-specific gender prediction algorithm? (Genni tool) based on local frequency statistics. A total of 35 out of 51 universities in our dataset, accounting for 53.28% of the 13,079 entries, did not include individual postdoc names. From the remaining 46.72% of entries: 41% are inferred men, 31.36% are inferred women, and 27.58% are unassigned (either because the algorithm did not produce a significant result, or due to names of particular national origins that allow “unisex” use).

#### Position titles

Postdoc titles are notoriously varied in the U.S. (McDowell, 2016; Schaller et al., 2017). Our dataset contained combinations of many of the same identifying words in specific position titles: assistant, teaching, clinical, scholar, senior, fellow, associate, researcher, faculty, intern, and trainee (*e.g.* Postdoctoral Research Fellow vs. Senior Postdoctoral Fellow), which were found in all but 395 of the 13,079 position titles. To examine possible associations between position titles and salaries, we searched for the presence of individual identifying words in each title entry. This process could result in a postdoc being represented more than once in the dataset if the position title contained more than one of these words (**Table 2**).

We utilized linear regression to determine the influence of the position title on salary. For the linear model, we transformed the categorical title description data into a sparse matrix where “1” represents the presence of the word and “0” represents its absence. This data configuration preserves the relationship between an individual’s salary and a specific combination of title words (*i.e.* each postdoc is counted only once). The model was calculated in R in the simple form:

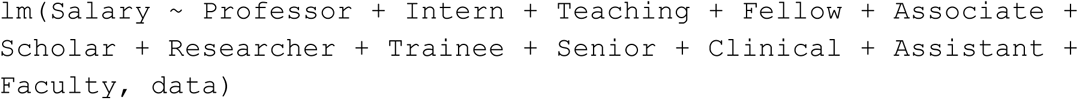

Or

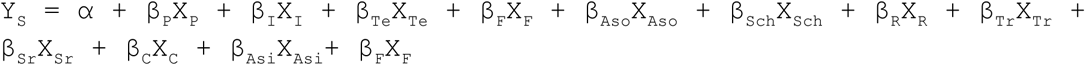

The results are presented in **Table 3**.

#### NIH funding

The total funds awarded by the NIH to each institution in 2017 were determined using NIH RePort (“NIH RePort”, n.d.) for the institutions where this data was available, which comprised 38 out of the 52 institutions in the dataset (**Table 4**). The standard deviation and coefficient of variation were calculated for the reported salaries from NIH-funded institutions. The coefficients of variation together with the total 2017 NIH award amount (USD) awarded to each institution were used to estimate the smoothed loess line and 95% confidence intervals of **Figure 4**.

## Results

### Initial studies and region-specific analyses

Our initial analysis of postdoc salary data for 13,079 postdocs across 52 institutions, estimated at 15-30% of the total U.S. postdoc workforce, is summarized in **Table 1**. The number of science, engineering and health postdocs in the NSF’s 2015 GSS data (National Science Foundation, n.d.) was used as an estimate of the approximate number of total postdoc salaries which we should have received from each institution. Salaries below the $23,660 legal minimum are assumed to be reporting errors (such as postdocs paid on direct fellowships where salary was provided directly from the funder, bypassing the institution’s payroll system), and were excluded from the subsequent analysis in **Table 1**. Institutional reporting of postdoc salaries was variable with respect to this issue. However, we independently verified that some institutions, such as the University of Washington, provided their actual annual postdoc salaries.

**Table 1 :**
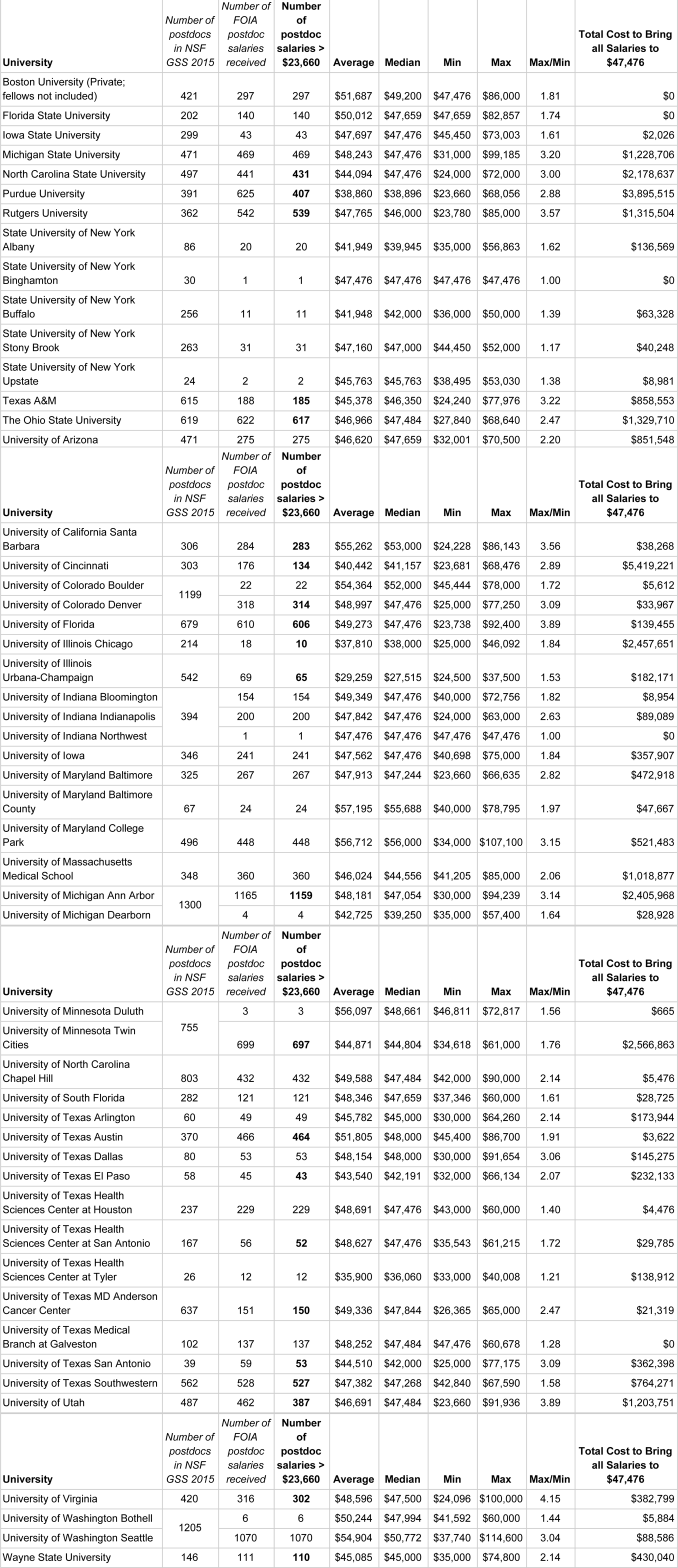
Comparison of postdoc salary information across U.S. institutions. The number of postdocs in the NSF’s GSS 2015 data is used as a comparison of the expected number of postdoc salaries that should have been received, to give us an indication of how much data may be missing from an institution. The number of postdoc salaries received that are greater than $23,660 are indicated and emboldened where they differ from the number of salaries received. This data was then used to calculate the average, median, maximum and minimum values, and to calculate the total cost to the institution of bringing all salaries up to the FLSA threshold salary that was proposed for December 1st 2016.

| **University** | *Number of*<br>*postdocs*<br>*in NSF*<br>*GSS 2015* | *Number of*<br>*FOIA*<br>*postdoc*<br>*salaries*<br>*received* | **Number**<br>**of**<br>**postdoc**<br>**salaries >**<br>**\$23,660** | **Average** | **Median** | **Min** | **Max** | **Max/Min** | **Total Cost to Bring**<br>**all Salaries to**<br>**\$47,476** |
| --- | --- | --- | --- | --- | --- | --- | --- | --- | --- |
| Boston University (Private;<br>fellows not included) | 421 | 297 | 297 | \$51,687 | \$49,200 | \$47,476 | \$86,000 | 1.81 | \$0 |
| Florida State University | 202 | 140 | 140 | \$50,012 | \$47,659 | \$47,659 | \$82,857 | 1.74 | \$0 |
| Iowa State University | 299 | 43 | 43 | \$47,697 | \$47,476 | \$45,450 | \$73,003 | 1.61 | \$2,026 |
| Michigan State University | 471 | 469 | 469 | \$48,243 | \$47,476 | \$31,000 | \$99,185 | 3.20 | \$1,228,706 |
| North Carolina State University | 497 | 441 | **431** | \$44,094 | \$47,476 | \$24,000 | \$72,000 | 3.00 | \$2,178,637 |
| Purdue University | 391 | 625 | **407** | \$38,860 | \$38,896 | \$23,660 | \$68,056 | 2.88 | \$3,895,515 |
| Rutgers University | 362 | 542 | **539** | \$47,765 | \$46,000 | \$23,780 | \$85,000 | 3.57 | \$1,315,504 |
| State University of New York<br>Albany | 86 | 20 | 20 | \$41,949 | \$39,945 | \$35,000 | \$56,863 | 1.62 | \$136,569 |
| State University of New York<br>Binghamton | 30 | 1 | 1 | \$47,476 | \$47,476 | \$47,476 | \$47,476 | 1.00 | \$0 |
| State University of New York<br>Buffalo | 256 | 11 | 11 | \$41,948 | \$42,000 | \$36,000 | \$50,000 | 1.39 | \$63,328 |
| State University of New York<br>Stony Brook | 263 | 31 | 31 | \$47,160 | \$47,000 | \$44,450 | \$52,000 | 1.17 | \$40,248 |
| State University of New York<br>Upstate | 24 | 2 | 2 | \$45,763 | \$45,763 | \$38,495 | \$53,030 | 1.38 | \$8,981 |
| Texas A&M | 615 | 188 | **185** | \$45,378 | \$46,350 | \$24,240 | \$77,976 | 3.22 | \$858,553 |
| The Ohio State University | 619 | 622 | **617** | \$46,966 | \$47,484 | \$27,840 | \$68,640 | 2.47 | \$1,329,710 |
| University of Arizona | 471 | 275 | 275 | \$46,620 | \$47,659 | \$32,001 | \$70,500 | 2.20 | \$851,548 |
| University of California Santa Barbara | 306 | 284 | **283** | \$55,262 | \$53,000 | \$24,228 | \$86,143 | 3.56 | \$38,268 |
| University of Cincinnati | 303 | 176 | **134** | \$40,442 | \$41,157 | \$23,681 | \$68,476 | 2.89 | \$5,419,221 |
| University of Colorado Boulder | 1199 | 22 | 22 | \$54,364 | \$52,000 | \$45,444 | \$78,000 | 1.72 | \$5,612 |
| University of Colorado Denver | | 318 | **314** | \$48,997 | \$47,476 | \$25,000 | \$77,250 | 3.09 | \$33,967 |
| University of Florida | 679 | 610 | **606** | \$49,273 | \$47,476 | \$23,738 | \$92,400 | 3.89 | \$139,455 |
| University of Illinois Chicago | 214 | 18 | **10** | \$37,810 | \$38,000 | \$25,000 | \$46,092 | 1.84 | \$2,457,651 |
| University of Illinois Urbana-Champaign | 542 | 69 | **65** | \$29,259 | \$27,515 | \$24,500 | \$37,500 | 1.53 | \$182,171 |
| University of Indiana Bloomington | 394 | 154 | 154 | \$49,349 | \$47,476 | \$40,000 | \$72,756 | 1.82 | \$8,954 |
| University of Indiana Indianapolis | | 200 | 200 | \$47,842 | \$47,476 | \$24,000 | \$63,000 | 2.63 | \$89,089 |
| University of Indiana Northwest | | 1 | 1 | \$47,476 | \$47,476 | \$47,476 | \$47,476 | 1.00 | \$0 |
| University of Iowa | 346 | 241 | 241 | \$47,562 | \$47,476 | \$40,698 | \$75,000 | 1.84 | \$357,907 |
| University of Maryland Baltimore | 325 | 267 | 267 | \$47,913 | \$47,244 | \$23,660 | \$66,635 | 2.82 | \$472,918 |
| University of Maryland Baltimore County | 67 | 24 | 24 | \$57,195 | \$55,688 | \$40,000 | \$78,795 | 1.97 | \$47,667 |
| University of Maryland College Park | 496 | 448 | 448 | \$56,712 | \$56,000 | \$34,000 | \$107,100 | 3.15 | \$521,483 |
| University of Massachusetts Medical School | 348 | 360 | 360 | \$46,024 | \$44,556 | \$41,205 | \$85,000 | 2.06 | \$1,018,877 |
| University of Michigan Ann Arbor | 1300 | 1165 | **1159** | \$48,181 | \$47,054 | \$30,000 | \$94,239 | 3.14 | \$2,405,968 |
| University of Michigan Dearborn | | 4 | 4 | \$42,725 | \$39,250 | \$35,000 | \$57,400 | 1.64 | \$28,928 |
| University of Minnesota Duluth | 755 | 3 | 3 | \$56,097 | \$48,661 | \$46,811 | \$72,817 | 1.56 | \$665 |
| University of Minnesota Twin Cities | | 699 | **697** | \$44,871 | \$44,804 | \$34,618 | \$61,000 | 1.76 | \$2,566,863 |
| University of North Carolina Chapel Hill | 803 | 432 | 432 | \$49,588 | \$47,484 | \$42,000 | \$90,000 | 2.14 | \$5,476 |
| University of South Florida | 282 | 121 | 121 | \$48,346 | \$47,659 | \$37,346 | \$60,000 | 1.61 | \$28,725 |
| University of Texas Arlington | 60 | 49 | 49 | \$45,782 | \$45,000 | \$30,000 | \$64,260 | 2.14 | \$173,944 |
| University of Texas Austin | 370 | 466 | **464** | \$51,805 | \$48,000 | \$45,400 | \$86,700 | 1.91 | \$3,622 |
| University of Texas Dallas | 80 | 53 | 53 | \$48,154 | \$48,000 | \$30,000 | \$91,654 | 3.06 | \$145,275 |
| University of Texas El Paso | 58 | 45 | **43** | \$43,540 | \$42,191 | \$32,000 | \$66,134 | 2.07 | \$232,133 |
| University of Texas Health Sciences Center at Houston | 237 | 229 | 229 | \$48,691 | \$47,476 | \$43,000 | \$60,000 | 1.40 | \$4,476 |
| University of Texas Health Sciences Center at San Antonio | 167 | 56 | **52** | \$48,627 | \$47,476 | \$35,543 | \$61,215 | 1.72 | \$29,785 |
| University of Texas Health Sciences Center at Tyler | 26 | 12 | 12 | \$35,900 | \$36,060 | \$33,000 | \$40,008 | 1.21 | \$138,912 |
| University of Texas MD Anderson Cancer Center | 637 | 151 | **150** | \$49,336 | \$47,844 | \$26,365 | \$65,000 | 2.47 | \$21,319 |
| University of Texas Medical Branch at Galveston | 102 | 137 | 137 | \$48,252 | \$47,484 | \$47,476 | \$60,678 | 1.28 | \$0 |
| University of Texas San Antonio | 39 | 59 | **53** | \$44,510 | \$42,000 | \$25,000 | \$77,175 | 3.09 | \$362,398 |
| University of Texas Southwestern | 562 | 528 | **527** | \$47,382 | \$47,268 | \$42,840 | \$67,590 | 1.58 | \$764,271 |
| University of Utah | 487 | 462 | **387** | \$46,691 | \$47,484 | \$23,660 | \$91,936 | 3.89 | \$1,203,751 |
| University of Virginia | 420 | 316 | **302** | \$48,596 | \$47,500 | \$24,096 | \$100,000 | 4.15 | \$382,799 |
| University of Washington Bothell | 1205 | 6 | 6 | \$50,244 | \$47,994 | \$41,592 | \$60,000 | 1.44 | \$5,884 |
| University of Washington Seattle | | 1070 | 1070 | \$54,904 | \$50,772 | \$37,740 | \$114,600 | 3.04 | \$88,586 |
| Wayne State University | 146 | 111 | **110** | \$45,085 | \$45,000 | \$35,000 | \$74,800 | 2.14 | \$430,040 |

The average, median, minimum and maximum salaries for each institution are listed in **Table 1**, the most useful data for comparisons between institutions being the median salary. The lowest median salary calculated from this dataset was University of Illinois Urbana-Champaign at $27,515, and the highest was University of Maryland College Park at $56,000. However the number of salaries received from the University of Illinois was much lower than expected, and the data at the lower end of the range could be an underestimate of the actual values due to differences in salary reporting, rather than actual lower salary values. Indeed, at time of writing, we had just received updated data from the University of Illinois which is currently being analyzed.

Following this initial analysis, we separated the aggregate data from 51 institutions into geographic regions (“Geography Atlas - Regions - Geography - U.S. Census Bureau”, n.d.) to show the distribution of postdoc numbers by institution that were grouped by U.S. region (**Figure 1**). The majority of institutions used in data collection were from the South and Midwest (20 and 16 institutions respectively). There was no difference in salary earned overall by postdocs between different regions (**Supplementary Figure 1**).

**Figure 1.**
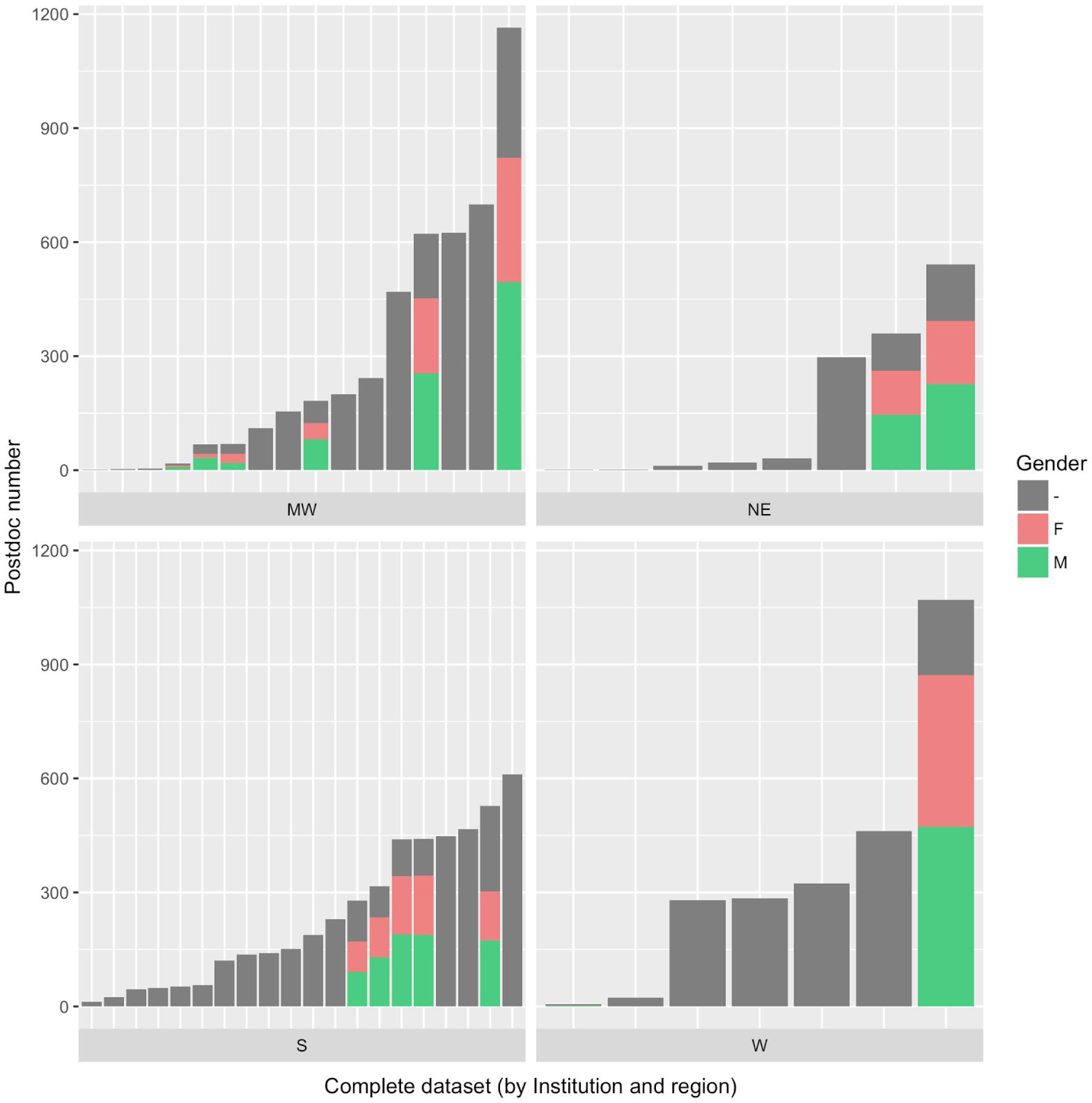
Number of postdocs at each institution by gender, and U.S. region. MW=Midwest, NE=Northeast, S=South, W=West. Each bar represents an institution with the specified geographic region. Gender was inferred from available names using the “Ethnea” algorithm. Lack of gender information (“-”, gray bar) signifies absence of name information or the inability of the algorithm to determine the person’s gender with confidence. Due to cultural norms in name-giving, names of particular national origins that allowed “unisex” use were not assigned a gender and are therefore depleted from the gendered population in this study.

### Salary Analysis by Discipline

We initially requested only information on postdoc salaries and job titles as part of our FOIA requests. However, for nearly half of the dataset, information was also received identifying postdoc department or school affiliation, which allowed us to evaluate whether lower salaries reported might be due to postdocs in “non-STEM” disciplines, who are assumed to be paid differently than those in STEM fields. Using this information, we classified each of the 2,883 postdocs from the 7 universities with available departmental information as “STEM”, “non-STEM”, or lacking sufficient information to unambiguously classify the department (**Supplementary Table 1**). Most salaries for “non-STEM” postdocs were in the same range as seen in our overall data, with most salaries in the $40,000-$50,000 range ((Woolston, 2017), **Supplementary Figure 2A**) and no differences in salaries were identified between STEM and “non-STEM” (**Supplementary Figure 2B**) although more data on postdoc affiliation, and perhaps even area of specific study, are needed to refine this analysis.

### Salary analysis by gender

In addition, we received names for 6,110 of the postdocs in our dataset. Although these entries did not include information on postdoc gender, we took advantage of the name information to estimate the effect of this variable on postdoc salaries. We inferred gender for each name using the Ethnea algorithm (Torvik and Agarwal, 2016) to 4,425 out of 6,110 entries with supplied names (16 universities, **Figure 1**). We estimate a male-to-female postdoc ratio of 1.3 across all universities. This is equivalent to 43% female postdocs in total and is in general agreement with previous studies (~40%, (Sheltzer and Smith, 2014)).

Subsequent analyses examined postdoc salaries by U.S. region and inferred gender (**Figure 2**), which comprise 891 male postdocs and 609 female postdocs in the Midwest, 372 male postdocs and 282 female postdocs in the Northeast, 771 male postdocs and 624 female postdocs in the South, and 471 male postdocs and 401 female postdocs in the West. These data show the average postdoc salary is significantly higher for male than female postdocs in universities across the Northeast and South ($1,666 and $1,944 higher, respectively; t-test, p-value 3×10^−3^ and 4×10^−6^ respectively).

**Figure 2.**
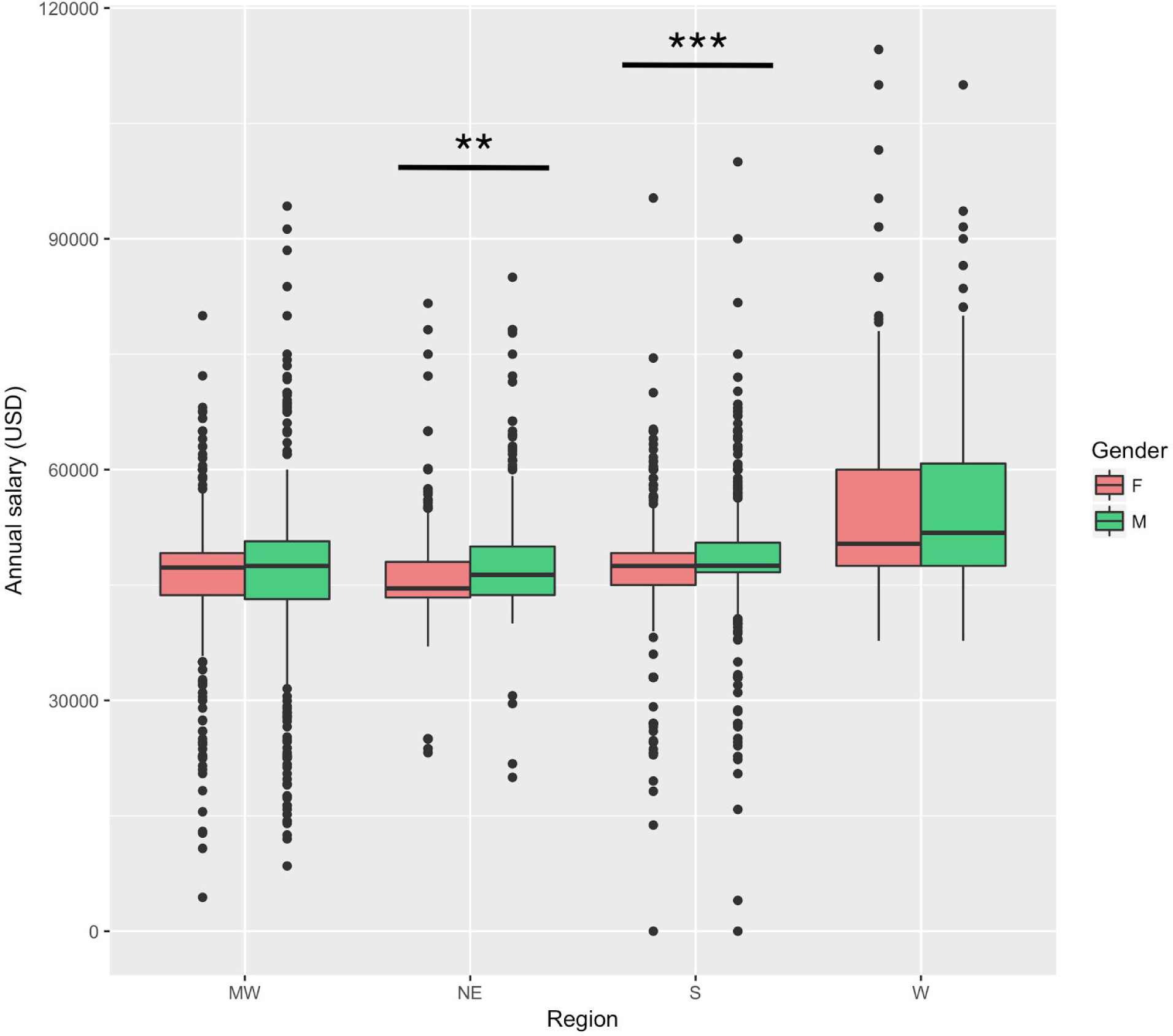
Postdoc salaries by U.S. region and inferred gender. MW=Midwest, NE=Northeast, S=South, W=West. Gender was inferred from available names using the “Ethnea” algorithm. Only the entries with available corresponding gender data (Figure 1) are shown. The means between genders are significantly different in the NE and S regions (t-test, ** p-value < 5 10^−3^, *** p-value < 5 10^−4^, respectively).

For universities in the West, the vast majority of salary data accompanied by gender assignment come from a single institution (see colored bars in **Figure 1**). Salaries from this institution do not seem to exhibit any systematic disparity between genders (average difference in salaries between male and female postdocs in the West: $406, **Figure 2**). The average difference in salaries between genders in the Midwest was also not significantly different ($764, **Figure 2**).

### Can varying postdoc titles predict salaries?

While we received names for only part of our dataset, we specifically requested postdoc position titles from all universities; all salary data we received included this information. Given the different identifying words in the position titles, as well as the multitude of existing postdoc titles (McDowell, 2016; Schaller et al., 2017) we were next interested to see whether actual position titles reflect different salary scales. For this analysis we mapped out the position titles assigned to all 13,079 postdocs from 51 institutions to 11 descriptive words (**Table 2**). As seen in **Figure 3**, we observed different salary ranges depending on the specific words used in each position title. As expected, the word “clinical” was associated with higher salaries. A higher salary was also observed for the description “faculty”, which was present in eight titles (**Table 2**), exemplifying the institutional variability in these titles.

**Figure 3.**
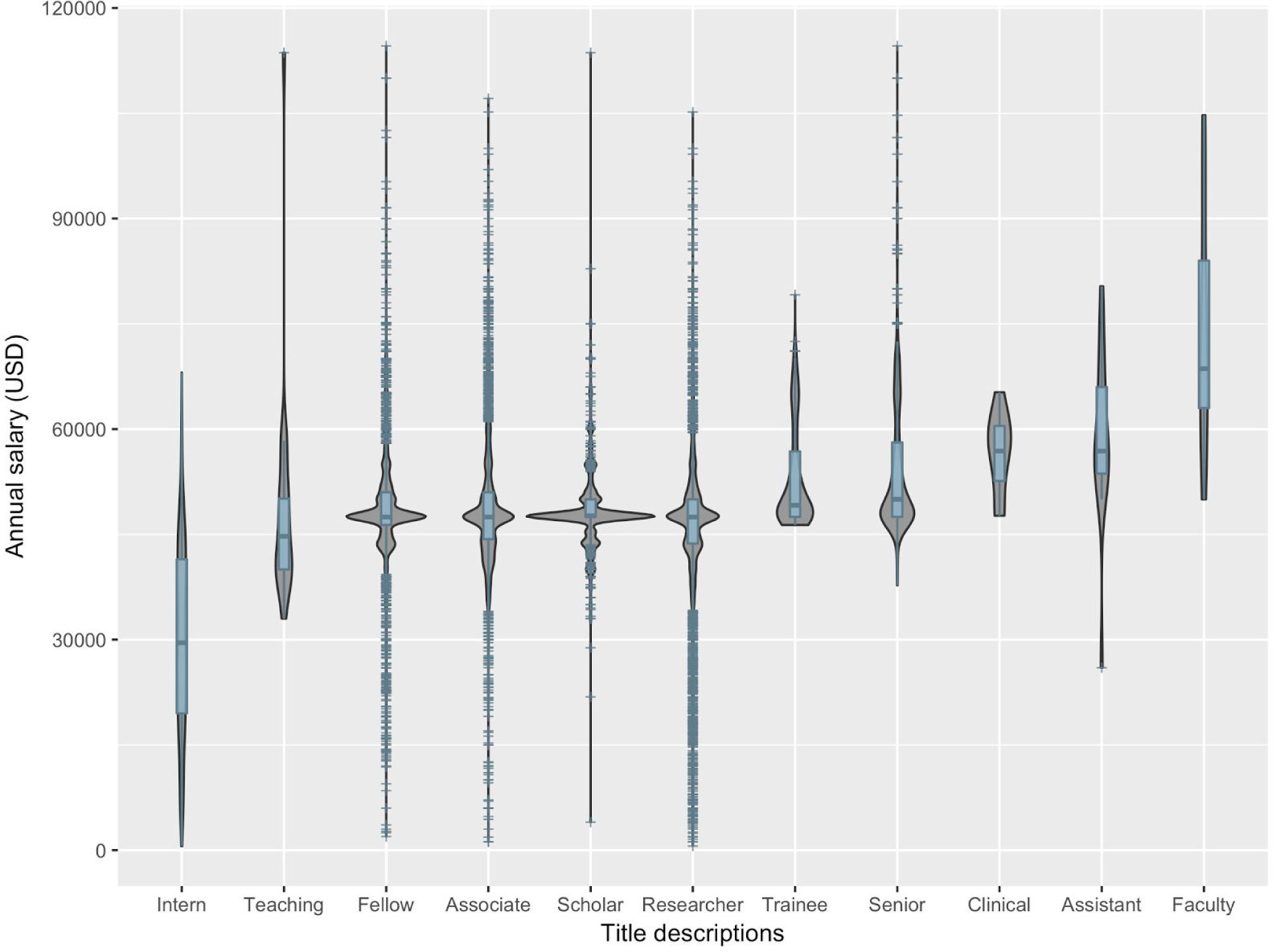
Salaries by position title description. The x-axis contains the descriptive words used in the various titles assigned to postdocs. As more than one word can be used in each title, the sum of the entries for each word exceeded the sum of postdoc numbers (Table 2, Methods).

**Table 2.** Position title descriptions by number of postdocs, and number of institutions using them.

| **Position title description** | **Number of postdocs** | **Number of institutions** |
| --- | --- | --- |
| **Intern** | 649 | 2 |
| **Teaching** | 18 | 1 |
| **Fellow** | 4643 | 29 |
| **Associate** | 5415 | 29 |
| **Scholar** | 792 | 5 |
| **Researcher** | 6336 | 28 |
| **Trainee** | 294 | 3 |
| **Senior** | 740 | 4 |
| **Clinical** | 17 | 3 |
| **Assistant** | 15 | 4 |
| **Faculty** | 8 | 1 |

Modelling the salaries as a linear function of the presence of each descriptive word in the job title allowed us to estimate the dollar impact of each individual term on the annual salary, without assuming a causal relationship (**Table 3**). The word “intern” appeared in 649 titles from 2 institutions (**Table 2**) and had a negative impact of $17,421 to a postdoc’s salary. On the other hand, the terms “trainee”, “senior”, and “clinical” added $1,801, $6,497, and $10,737 to the salary, respectively (**Table 3**). Of the model terms that did not significantly affect the outcome (**Table 3** : p-value, non-bolded text), the words “teaching” and “assistant” are underrepresented in our data (**Table 2**) and more information is needed to conclude whether they may have an effect on salary.

**Table 3.** The effect of position title description words on salaries. (p-values below 0.005 are denoted with bold text).

| Term | Value (\$) | error | p-value |
| --- | --- | --- | --- |
| Intern | -17,421 | 435 | **0.000** |
| Teaching | 889 | 2,206 | 0.687 |
| Fellow | -641 | 284 | 0.024 |
| Associate | 614 | 272 | 0.024 |
| Scholar | 514 | 409 | 0.209 |
| Researcher | -185 | 178 | 0.306 |
| Trainee | 1,802 | 589 | **0.002** |
| Senior | 6,479 | 390 | **0.000** |
| Clinical | 10,737 | 2,506 | **0.000** |
| Assistant | -5.008 | 3,190 | 0.420 |
| Faculty | 26,688 | 4,043 | **0.000** |

### Examining the relationship between salary (per-postdoc) and NIH funding by institution

We expected institutions receiving more NIH funding to more readily be able to raise postdoc salaries, or be in a better position to maintain high postdoc salaries. To investigate this question, we examined the relationship between total NIH funding received by an institution in 2017 and the salaries of postdocs employed there. We found that the number of postdocs tends to decrease at institutions with smaller or fewer NIH grants received in 2017 (**Table 4**, pearson correlation: 0.76). However, institutions with an average or relatively low amount of NIH funding maintain a large postdoctoral population.

**Table 4.** Institutions ranked by total 2017 NIH funding with number of postdoc salaries. “ % grant $ in set” denotes what percentage of the total funding given to the 38 institutions is received by that institution.

| NIH awards rank | % grant \$ in set | Postdoc # | | NIH awards rank | % grant \$ in set | Postdoc # |
| --- | --- | --- | --- | --- | --- | --- |
| 1 | 13.17 | 1165 |  | 20 | 1.60 | 183 |
| 2 | 10.62 | 440 |  | 21 | 1.58 | 56 |
| 3 | 5.58 | 324 |  | 22 | 1.53 | 18 |
| 4 | 4.60 | 528 |  | 23 | 1.51 | 469 |
| 5 | 4.39 | 622 |  | 24 | 1.42 | 111 |
| 6 | 4.31 | 297 |  | 25 | 1.40 | 188 |
| 7 | 4.26 | 462 |  | 26 | 1.21 | 448 |
| 8 | 4.16 | 610 |  | 27 | 1.16 | 625 |
| 9 | 4.11 | 278 |  | 28 | 1.09 | 23 |
| 10 | 3.96 | 360 |  | 29 | 0.93 | 140 |
| 11 | 3.72 | 243 |  | 30 | 0.60 | 441 |
| 12 | 3.72 | 151 |  | 31 | 0.43 | 53 |
| 13 | 3.60 | 316 |  | 32 | 0.40 | 284 |
| 14 | 2.92 | 121 |  | 33 | 0.37 | 68 |
| 15 | 2.44 | 280 |  | 34 | 0.35 | 45 |
| 16 | 2.38 | 229 |  | 35 | 0.17 | 1 |
| 17 | 2.32 | 466 |  | 36 | 0.14 | 49 |
| 18 | 2.03 | 137 |  | 37 | 0.12 | 12 |
| 19 | 1.66 | 31 |  | 38 | 0.02 | 3 |

Plotting the salary distributions per institution, ranked in descending order by the amount of NIH funding received (**Figure 4A**), demonstrates that regardless of NIH award rank, the median postdoc salary is centered around the NRSA minimum of $47,484 at most institutions in this dataset. However, this is more variable for some institutions receiving less NIH funding, where the salary distribution is less homogeneously centered around the population’s mean. This variability cannot be readily explained by noise, but may be due to lower numbers of postdocs at these institutions (**Table 4**). We also examined the standard deviation (which reflects both sample size and variance) of postdoc salaries for each institution and calculated the salary coefficient of variation (**Figure 4B**). Plotted against each institution in Figure 4A, this metric confirms the variability observed in salaries per institution.

**Figure 4.**
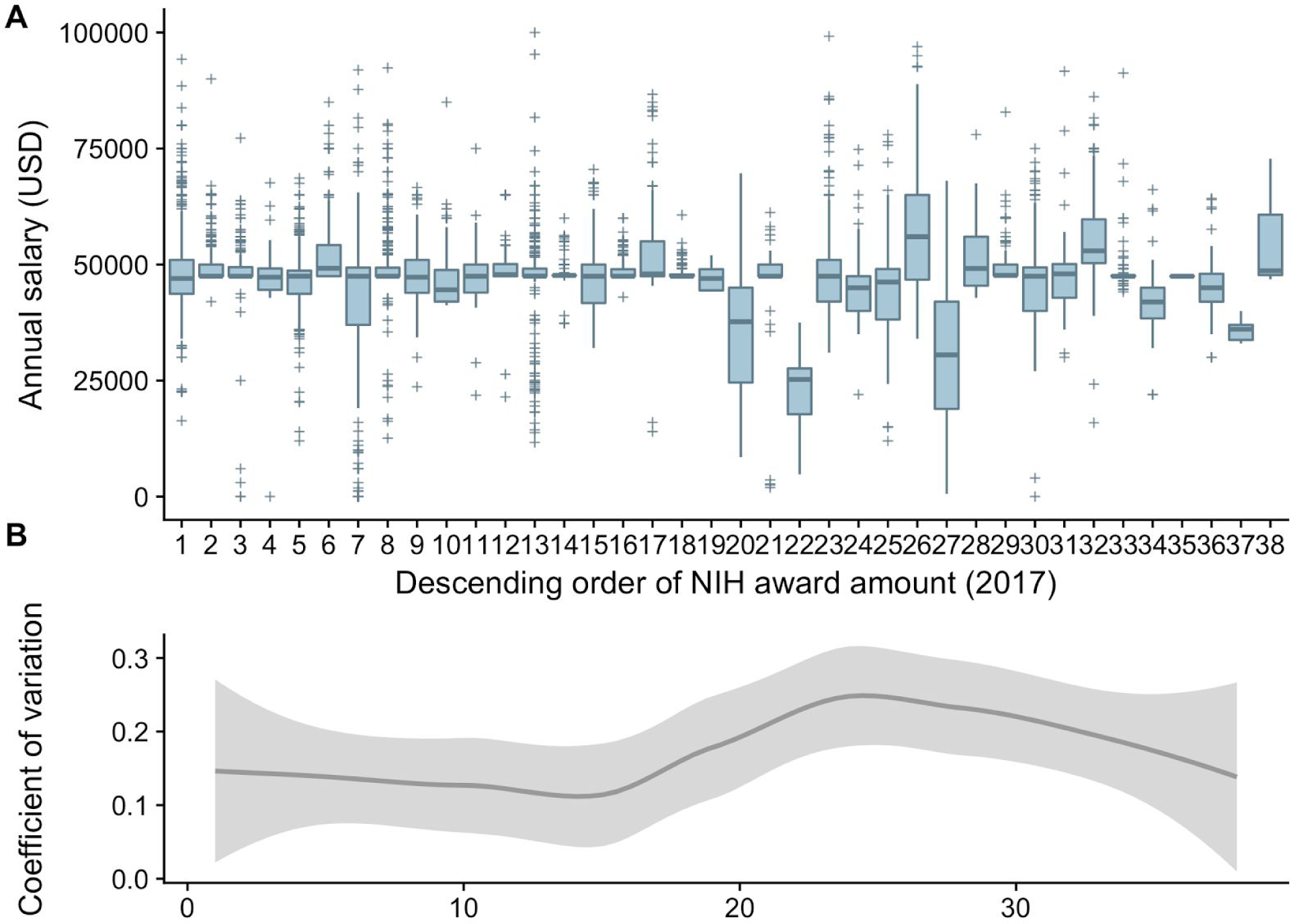
Postdoc salaries as a function of total 2017 NIH funding. Each institution is depicted numerically in the x-axis, in descending order of the total amount of 2017 NIH grant funding awarded (1=best funded, 38=least funded). Lower NIH funding levels are associated with higher postdoc salary variability. **(A)** Boxplots of salary distribution per institution. **(B)** Smoothed (loess) line of the coefficient of variation (standard deviation/mean) of the salaries in each Institution. The grey area denotes 95% confidence intervals.

## Discussion

The data we have gathered on over 13,000 postdoctoral salaries and job titles across 52 U.S. academic institutions aimed to increase transparency about the landscape of postdoctoral salaries in the U.S. In addition to these data, we have published monthly blog posts with data from each institution on our website resource (“Investigating Postdoc Salaries | Future of Research”, n.d.) every day for the entire month of November 2017. This exercise has resulted in open and informal discussions of caveats and limitations of these data, and has allowed a comparison of data between institutions.

This is a work in progress, since in preparing this manuscript, even on the day of submission, we continue to receive clarifications, updates and additions to the data received. **We are in the process of analyzing these submitted datasets, and welcome community suggestions and any additional data from both public and private institutions in order to improve this work in progress and increase transparency**. We view this as a long-term project in concert with the efforts of other groups to both clarify postdoc titles and harmonize postdoc administration in general (Schaller et al., 2017). In particular, we have used the aggregate data collected here using standardized methodologies and are currently discussing future directions for this project, which include using data which has been provided voluntarily or clarified for us by other groups, as opposed to or in addition to data with recognized flaws, but which is collected in a standardized manner.

As seen from **Table 1** and discussed in (Woolston, 2017), median postdoc salaries across U.S. institutions, and a large proportion of individual salaries, cluster around the proposed annual salary levels according to the FLSA threshold for overtime exemption as updated in 2016 ($47,476) (Bankston and McDowell, 2016), and the NIH NRSA Year 0 salary for FY 2017 ($47,484, (National Institutes of Health, 2016). Thus, approximately 22% of all salaries are within a $25 range of those benchmarks ($47,475-$47,500). While we do not have postdoc salary data from prior years to compare to, those new FLSA and NIH salary levels appear to have had a major impact on determining the standard for postdoctoral salaries in the corresponding financial year. However, the NIH stipend levels only apply to postdocs on NIH NRSA funding mechanisms, and do not apply to all postdocs employed at U.S. institutions, including those on NIH research grants such as R01s. While there is no federal mandate for annual salaries to be above $23,660 for most postdocs, it is striking the potential effect a now-defunct federal mandate, and benchmarking set by NIH, may have had on postdoc salary norms.

Delving more deeply into variability in our dataset, it is possible that lower salaries may be due to means by which postdoc compensation is recorded, tracked and reported by individual institutions. However these lower salaries may not be entirely attributed to postdocs in non-STEM fields (**Supplementary Figure 2**). Likewise, higher salaries in this dataset may be due to clinical research fellows, “career postdocs”, or simply inappropriate uses of postdoc titles for positions that are not actually postdocs, such as administrative staff and faculty. These higher-paid individuals may also not be primarily performing research in a short-term position, and may be conducting different activities to postdocs. However, these higher salaries may also reflect pay of traditional research postdocs, who may be compensated far better than most of their postdoctoral peers, perhaps due to experience, cost of living, or other factors. Therefore, negotiation for higher salary may be an option utilized successfully by a subset of postdocs.

A recently-published Life Science Salary Survey encompassing more than 2,500 life science professionals from around the world showed very little difference between the salaries of men and women in U.S. academia, and suggested that salary differences disappeared with more aggregated data (Mika, 2017). In relation to this topic, we find significant differences between the salaries of male and female postdocs in our sample in the NE and S regions, but not in the MW and W regions of the U.S. (**Figure 2**). This raises the question of how multiple regional variables within institutions in the U.S. and across the world may affect postdoc salaries. While postdoc salaries nationally may show general equity, future work should consider whether potential differences on local or regional levels may be masked by overall national trends.

We were also intrigued to find that certain position titles from our dataset could be correlated with salary data, which may indicate compliance to a certain regulation. With respect to specific titles, “trainee” could potentially be associated more closely with postdocs on NRSA fellowships, although this remains to be verified on a larger scale. Likely the word “clinical” would suggest higher salaries, whereas the “teaching” group data is slightly surprising given that teaching postdocs are not subject to the FLSA, meaning their values may have been expected to be lower than we find in this dataset. However, the teaching population may also contain postdocs on IRACDA fellowships (who perform research 75% of the time), and a more thorough analysis of this population is needed. Finally, the “intern” group (observed at only two universities, North Carolina State and Purdue) is an interesting one, given that postdocs are not typically associated with this word, and this could significantly affect their pay level.

For the purposes of the manuscript, given the multitude of existing titles, we opted to use the term “postdoc” for consistency. However, we call for a universal system by which institutions may better document and report postdoc numbers and salaries across the U.S., enabling improved reporting on specific aspects of the postdoctoral position, including career outcomes tracking of the biomedical workforce encompassing both graduate students and postdocs (Polka et al., 2015). This type of information would allow early career researchers to make informed career decisions about whether to pursue postdoctoral training and what aspects (including financial) they should consider when making these decisions. A unified system for tracking postdocs could also extend to other aspects of the postdoctoral position, such as career-specific achievement metrics, allowing for even greater transparency on multiple aspects of academia. Finally, complete datasets on particular aspects of the postdoctoral experience may aid in better training and support for postdocs both at the institutional and national level.

### Limitations of this work

Our findings indicate that several variables contribute to the ability of the scientific community to be aware of how much U.S. postdocs are actually being paid. The largest barrier for our study is the quality and completeness of available institutional data, and the willingness of institutions to make their postdoc salary data public. Institutional barriers to maintaining and/or providing these data may be related to nuances of institution-specific record keeping, the existence of a postdoc office at the time of data collection, or staff turnover in the individuals who have managed these data over time. Other limitations of this work include the fact that not all institutions reported salaries for all postdocs at their institution, and the majority of these data were obtained from public institutions.

In order to truly increase transparency on postdoc salaries across U.S. institutions, future work should aim to include data from both public and private institutions. It should also work to develop standardized data collection methods to enable analyses similar to those completed in this study. Due to these limitations, our data collection efforts constitute only a small portion (~13,000 postdocs, or roughly 15-30% of the postdoctoral workforce) of the total salary data available nationwide at universities, encompassing anywhere between ~50,000-100,000 postdocs (Biomedical Research Workforce Working Group, 2012; National Science Foundation, n.d.; Pickett et al., 2017; Schaller et al., 2017). Finally, our FOIA requests did not ask for detailed demographic or other descriptive data (e.g. postdoc department or discipline), which further limited our ability to characterize the effect of additional variables for postdocs receiving these salaries.

Using postdoc names supplied in this dataset together with nearly half of the analyzed salary data, we hoped to analyze not only the effect of gender, but also that of race, ethnicity, and national status, on postdoc salaries. However, without reported data on these demographics, and relying purely on predictive algorithms, we encountered limitations on the ability to pursue such an analysis. We strongly encourage others asking such questions in this space to consider whether there are differences between foreign-born researchers (who comprise approximately two-thirds of the U.S. biomedical postdoc population (Garrison et al., 2005), and more than half of the U.S. biomedical workforce as a whole) as well as researchers from underrepresented populations, to examine whether differences in postdoc salaries at institutions arise due to these variables (Heggeness et al., 2016, 2017).

## Future directions

As we plan to repeat this data collection exercise in the future to attempt a more longitudinal analysis, we may consider expanding data requests, as certain data were received automatically and may be readily available. In addition, the utility of using FOIA requests to gather this data offers insight into the capabilities of differing institutions to report data in this fashion, presumably requiring staff unfamiliar with postdocs to communicate with other branches of institutional administration in order to obtain this data. It was encouraging to see that some institutions were able to provide the exact data requested, and in the expected quantity, which indicates that at least some institutions are able to accurately track their postdocs. Ultimately, this may be an indicator of their institutional commitment to the development of postdoctoral researchers. We hope to see more institutions show this type of dedication in the future.

## Author contributions

G.S.M. conducted the FOIA requests, collected the data and performed preliminary analyses. R.A. analyzed the data, made the figures and wrote relevant paper sections. R.A., A.B., M.C., C.N. and G.S.M. contributed to writing and editing the paper. The final version was approved by all authors.

## Acknowledgements

G.S.M. is supported by the Open Philanthropy Project and, where needed, this support was used also to pay for FOIA requests. Open Philanthropy had no role in the design or undertaking of this work. We thank FoR advisory board member, Sarah Hokanson, for providing postdoc salary data for the only private institution in this data set (Boston University). We are also very grateful to those at institutions who have sent us data or discussed this data collection effort with us, and clarified issues that we encountered with the quality of the data.

## Supplementary Figures

**Supplementary Figure 1.**
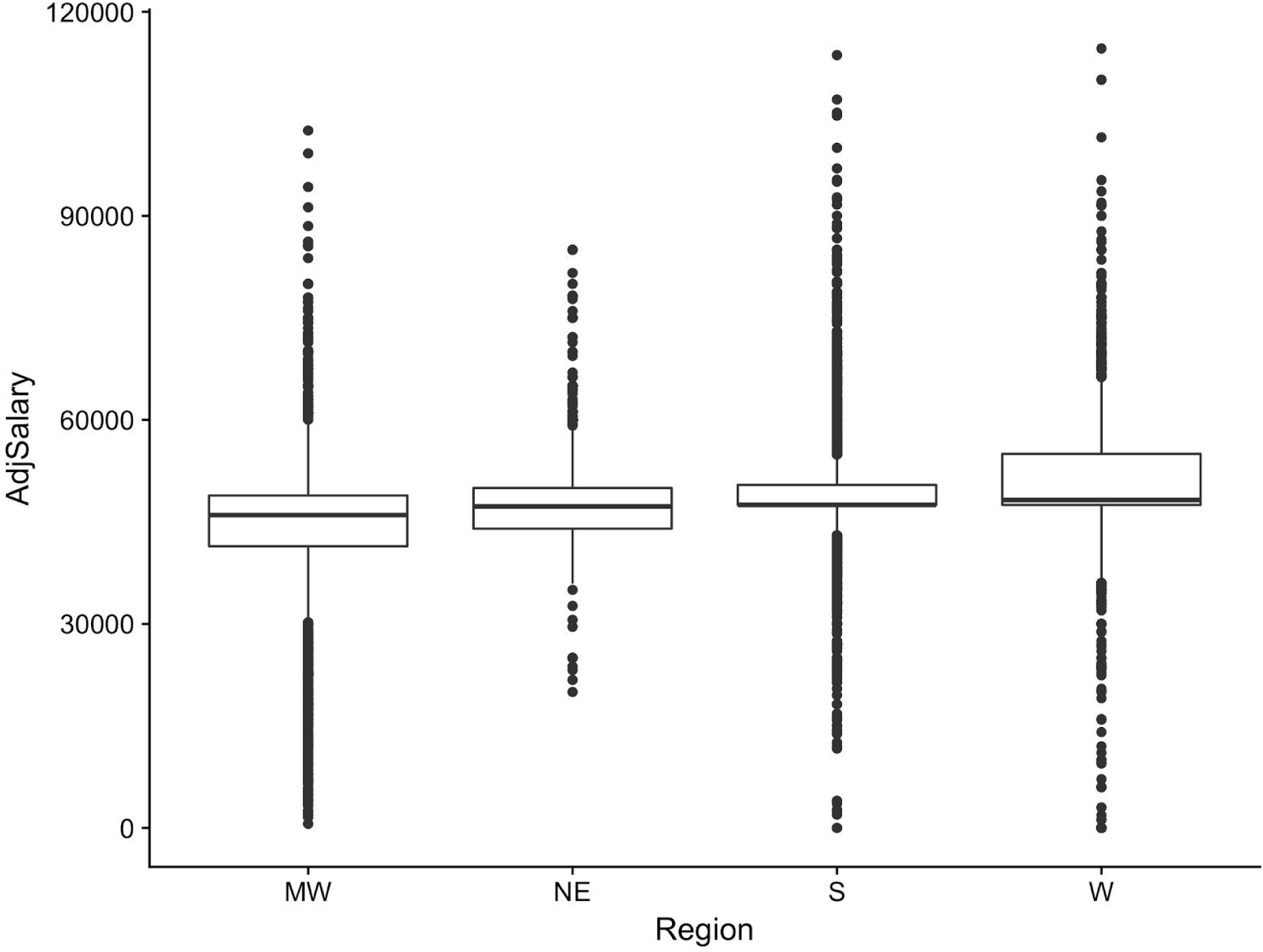
Postdoc salary ranges by geographic location of institutions. MW=Midwest, NE=Northeast, S=South, W=West.

**Supplementary Figure 2.**
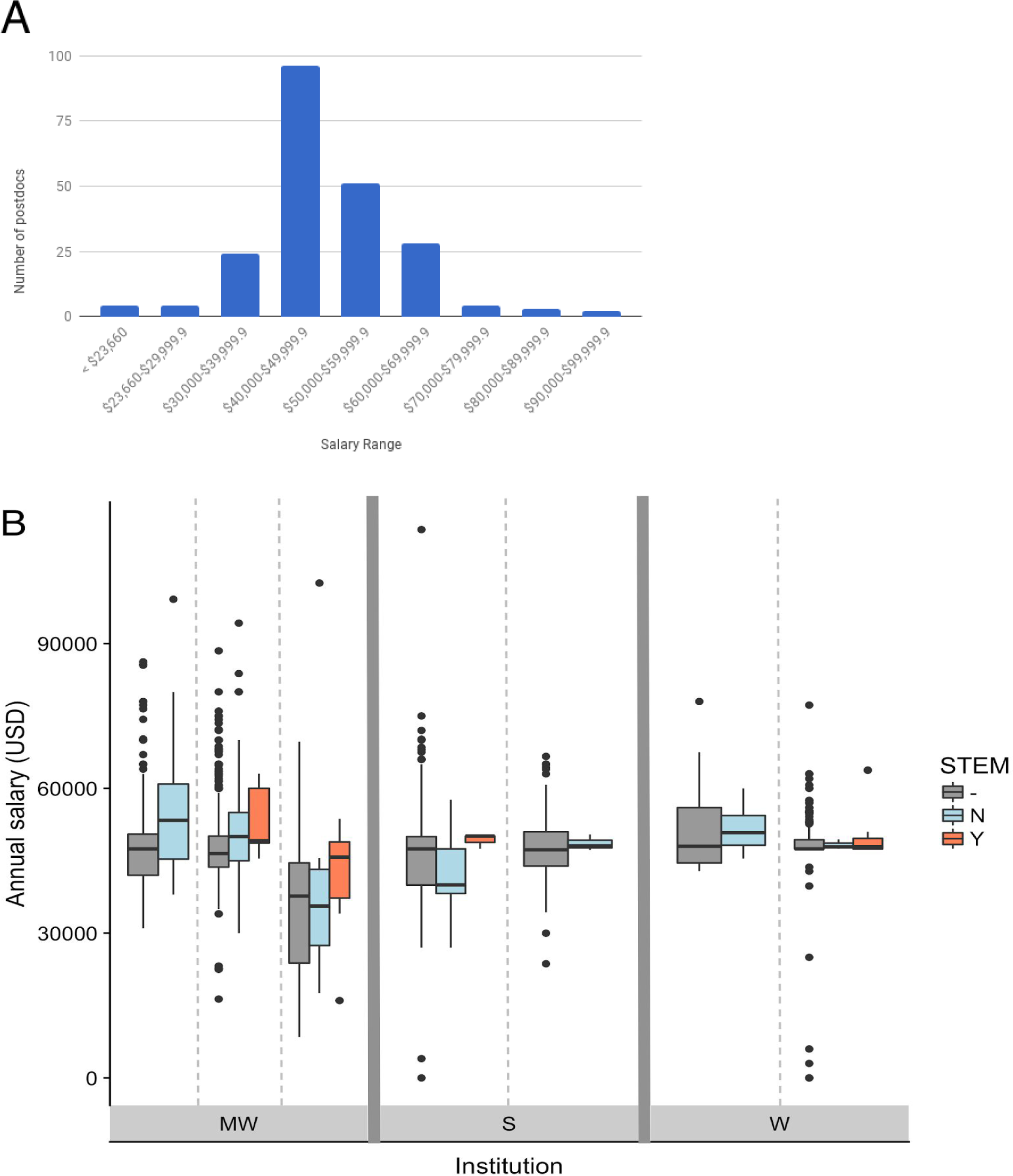
Annual salaries for non-STEM postdocs. **(A)** Distribution of salary ranges in non-STEM disciplines (n = 179 postdocs). **(B)** Comparison of salary ranges between postdocs in STEM and non-STEM research disciplines for 6 institutes with the relevant departmental information organized by region. MW=Midwest, S=South, W=West (no relevant departmental data was available from the N region). Legend: N: non-STEM, Y: STEM,“-” : entries from the same institution without departmental information, or without sufficient information to unambiguously classify the department.

**Supplementary Table 1.** Disciplines/departments employing non-STEM postdocs in this analysis.

|  |  |  |
| --- | --- | --- |
| Child Behavioral Health | Counseling Services | Political Studies |
| Depression Center | Public Policy | Health Behavior & Education |
| Law | Economics | English Language & Literature |
| History | History of Art | Humanities Institute |
| Chinese Studies | Korean Studies | Judaic Studies |
| Linguistics | Near Eastern Studies | Philosophy |
| Psychology | Nutritional Sciences | Population Studies |
| Education | Social Work | Youth & Social Issues |
| Urban Planning | Criminal Justice | Communication Sciences & Disorders |
| Marketing | Geography | Agricultural Food/Resource Economics |
| Communication | Media & Information | Food Science and Human Nutrition |
| Political Science | Teacher Education | Human Development and Family Studies |
| English | Religious Studies | Foreign Languages and Literature |
| Sociology | Anthropology | Parks, Recreation, Tourism Management |

